# What can Y-DNA analysis reveal about the surname Hay?

**DOI:** 10.1101/2025.07.09.664039

**Authors:** Philip Stead, Penelope R Haddrill, Alasdair F Macdonald

## Abstract

The family name Hay (plus associated spelling variants) is a prominent Anglo-Norman-in-origin surname that has been well-documented as a Scottish noble lineage since the 12th century CE. Their historical significance, linked to the rise of the Anglo-Norman era (1093-1286 CE) in Scotland, and the historical complexities of surname adoption post-Norman conquest of England, justifies the need for a comprehensive understanding of the genetic history of the Hay noble lineage. This study focuses on examining the patterns of paternal inheritance in lineages with the Hay surname. We conducted a comprehensive analysis of Y-chromosome data that is publicly available on the Family Tree DNA (FTDNA) platform, and specific FTDNA surname projects, as well as looking in more detail at three well-documented male-line descendants of William II de la HAYA, 1^st^ of Erroll, (d. 1201) that have been verified to a high degree of confidence. Our results reveal that all descendants of William II de la HAYA, 1^st^ of Erroll, (d. 1201) derive from the multigenerational Y-SNPs R1a-YP6500 (plus equivalent SNPs BY33394 / FT2017) and R1a-FTT161. Furthermore, subclades of R1a-FTT161 have been identified that confirm direct male-line descent from two of William II de la HAYA’s sons. Subclade R1a-BY199342 (plus equivalents) confirms direct male-line descent from David de la HAYA, 2^nd^ of Erroll, (d. 1241), and subclade R1a-FTA7312 confirms direct male line decent from Robert de la HAYA of Erroll. The result also confirms that the Hay noble lineage shares the Y-SNP R1a-YP4138 (estimated to have occurred 832 CE) with several non-Hay testers that have surnames of Norman origin, therefore, providing further evidence to support the Norman origin hypothesis for these surnames. In addition to the identification of multigenerational Y-SNPs associated to documented Hay noblemen, this study has observed significant Y-DNA haplogroup diversity among males with the surname Hay (plus associated spelling variants: Hays, Haye, Hayes, Hey and Haya). Our results show that only 22% of the men sampled (n=109) with the surname Hay (plus associated spelling variation) are descended from the 12th century progenitor of the noble Hay lineage of Scotland. Therefore, confirming that a significant proportion of males with the surname Hay do not descend from the noble progenitor of the surname.

## Introduction

The lands of northern Britain that are now referred to as Scotland have not always been a unified realm. Between the early seventh century until the mid-ninth century, northern Britain was separated into four distinct kingdoms (1). The kingdom of Strathclyde was ruled by the Britons and incorporated the southwest region of modern-day Scotland; however, a large portion of this kingdom was incorporated into the Anglo-Saxon kingdom of Northumbria by the early eighth century which included the Lothians and the southeast border regions (1). The lands that primarily encompassed the northeastern third of Scotland, stretching roughly from the Firth of Forth in the south to Caithness and Orkney in the north was considered the kingdom of Pictavia, and ruled by the Picts (1). The kingdom of Dál Riata was ruled by the Scots, and their kingdom incorporated the western lands north of Loch Lomond and the Hebrides Islands (1).

The four kingdoms remained relatively stable until a significant event took place in 844 CE under the leadership of King Kenneth I (Cináed mac Alpin, died in 858 CE) who successfully unified the kingdom of Dál Riata and Pictavia into a single kingdom often referred to as Alba (2). Although the formation of the kingdom of Alba is generally considered the birth of Scotland, Broun (2015) argues that it is a significant oversight and oversimplification of the historical events that took place (3). Broun (1997) argues that it would not be until the thirteenth century, during the Anglo-Norman era, when the term Scotia (Scotland) was generally used when referring to the whole kingdom (4).

## The Anglo-Norman era in Scotland (1093-1286 CE)

The revolutionary change in the governance of Scotland, termed the ‘Anglo-Norman era’, was mainly facilitated by King David I of Scotland during his reign (c. 1124-1153 CE) (5). However, the start of Norman influence in Scotland started with the Treaty of Abernethy, where King Malcolm III of Scotland swore fealty to King William I of England in 1072 CE (6). King William I of England died on 9 September 1087 CE, succeeded by his son William II (commonly referred to as William Rufus). Malcolm III of Scotland went on to break the terms of the Treaty of Abernethy by invading northern England several times in the early 1090s CE (6). On the 13 November 1093 CE, King Malcolm III of Scotland invaded Berwick in northern England with his son Edward (6). King Malcolm III of Scotland advanced to Alnwick and was ambushed by English forces lead by Robert de Mowbray, Earl of Northumbria. Both King Malcolm III of Scotland and his son Edward were killed, leaving the Scottish forces without leadership, forcing their withdrawal back to the safety of Scotland, and facilitating the start of the Anglo-Norman era (c. 1093-1286 CE) (6).

King David I of Scotland (1084-1153 CE) had a new vision for Scotland, an aspiration to replicate the Norman feudal system imposed in England by King William I of England (c. 1066 CE) (5). To achieve his goal of adopting feudal tenure during his reign (c. 1124-1154 CE), King David I of Scotland attracted Anglo-Normans (people of higher social status of Anglo-Norman, French, and Flemish origin) to Scotland by offering Scottish lands, royal official appointments, noble titles, and knighthoods in return for feudal services (5). This invitation radically changed the governance of Scotland and impacted on the demographics of the Scottish population to an unknown degree (7).

## The documented origins of the Hay surname and the earliest Hay progenitor in Scotland

Among the foreign Anglo-Norman settlers of Scotland was the progenitor of the Scottish noble lineage Hay of Erroll, described as William de la HAYA II, 1^st^ of Erroll, (d.1201 CE) (8). William de la HAYA II, 1^st^ of Erroll was a Norman knight who is documented as cupbearer (butler) of King William the Lion of Scotland who succeeded King Malcolm IV on 9 December 1165 CE (9). William de la HAYA II, 1^st^ of Erroll (d. 1201 CE) is believed to be the son of William de la HAYA I (born in Cotentin, Normandy) and Julianna de Soulis; sister of Ranulf de Soulis, Lord Liddesdale (9), butler of King Malcolm IV, (c. 1153-1164 CE) (9–11). Black (1946) argues that there are several village names starting with ‘La Haye [haie/haia]’ in Normandy so one of these villages would be a good candidate for the origins of the surname Haya (12) (e.g., La Haye-du-Puits or La Haye Bellefond, both in the Soules region). Ritchie (1953) also supports La Haye-du-Puits as a possible origin of the Hay noble lineage of Scotland; however, he offers La Haye-de-Herce and La Haye Malherbe as potential alternatives (13). Moncrieffe (2010) argues that there is no doubt that their origins were from Haye-Hue, (Haia-Hugonis, now La Haye-Bellefond) because the La Haye-Hue of Normandy bear the same three escutcheons coat of arms used by the Hays of Erroll, and they also marched with de Soules [Soulis] near St Lo in the Cotentin peninsular of Normandy (14).

Barrow (1973) also supports Haye-Hue (La Haye-Bellefond) as the place of origin of the Hay noble lineage (15). The documented links between La Haye-Hue of Normandy, de Soulis (Normandy and Liddesdale), and Hay of Erroll are not likely to be coincidental. The Hay noble lineage firmly established their presence in Scotland, taking control of significant lands located in the Scottish Lowlands (16), and playing a substantial role in Scottish history thereafter. For example, Sir Gilbert de la HAY (d. 1333) was the fifth feudal Baron of Errol, Lord High Constable of Scotland, and companion of King Robert the Bruce of Scotland. Sir Gilbert de la HAY was one of the Scottish noblemen that signed the Declaration of Arbroath on 6 April 1320 CE (17).

The evidence that links William de la HAYA I to Normandy is that ‘la Haya’ is likely a Norman toponymic name, deriving its origin from a place called ‘la Haye/Haie’ in Normandy. McClure (2015, 2020) argue the case that surnames such as ‘de la Haye’ and ‘de la Mare’ often denote high ranking people from estates in Normandy (e.g., a notable individual from la Haie ‘the enclosure’ or la Mare ‘the pool’) (18, 19). However, McClure is also appropriately cautious in linking these locations to one specific origin or family (19). There are several villages in Normandy that have such place names, and Norman placenames were also introduced to Britain via the linguistic legacy of the Norman Conquest of 1066 CE. When William the Conqueror and his followers introduced Norman French culture, governance, and language to Britain, placenames often reflected the new Norman elite’s language, ownership, heritage, or religious affiliations, and were either imposed on new settlements or modified from existing Anglo-Saxon placenames. A good example is the placename Bernard Castle in County Durham, a place deriving its name from Bernard Baliol I (died before 1167 CE), a baron of Norman descent (20). Other examples include Norman French placenames derived from descriptive words such as Belvoir (bel voir) in Leicestershire, meaning ‘beautiful view’ (21) and Richmond in North Yorkshire that is derived from ‘riche mont’, meaning ‘strong hill’ (21, 22). Therefore, there is a strong possibility that many unrelated people adopted these surnames independently. Durie (2022) also highlights the complexities involved in identifying surname origins in his essay on clans, families and kinship structures in Scotland (23). He refers to the Hay noble lineage as being among the most influential families in Scotland; however, he does not discuss their potential origins prior to Scotland (23). Durie also highlights that having a surname that is specifically associated to nobility does not guarantee direct paternal descent from that noble lineage (23).

## Surname changes

Surname changes did historically take place with no documentational evidence to explain why/how the change occurred. Instances of surname changes were often the result of adoptions, divorce, out-of-wedlock conceptions, and extra-marital conceptions, so these situations would often remain undocumented (24). The genealogical term for these occurrences is a ‘non-paternity event’ (NPE) or a ‘false paternal event’ (FPE) (24), while Larmuseau et al (2017) uses the term ‘extra pair paternity’ (EPP) (25). Avni et al, (2023) suggested that the chances of an EPP taking place was between 1-5% in each generation, with the chances of discovering an EPP increasing with each successive generation (e.g., there is a 10-50% chance of discovering an EPP over ten-generations) (24). Moreover, King et al, (2014) revealed multiple instances of non-paternity in King Richard III’s family tree through the DNA analysis of material collected from his skeletal remains (26).

Traditional genealogical research heavily relies on documentary evidence to prove genealogies (27, 28). The documentary evidence strongly suggests that William de la HAYA II has direct paternal descendants surviving to the present (8, 29). However, traditional genealogical evidence alone is limited because it cannot identify an EPP occurring in any of the intervening generations of a pedigree (23) unless they were documented. Based on the evidence provided by Avni et al (2023), there is a good chance that someone with the surname Hay does not descend from William de la HAYA II (24). Furthermore, to the best of the authors’ knowledge, William de la HAYA I’s proposed link to Normandy is only supported by limited toponymic evidence so the pursuit of further evidence to explore his origins is well justified.

### Genetic Genealogy

The advancements in DNA sequencing technology, and the development of haplotype-based methods to detect population substructure have proven to provide insight into the degree of shared ancestry between populations (30–33). A recent study by Morez et al (2023) implemented FineSTRUCTURE clustering analysis and Identity-By-Descent (IBD) on imputed diploid dataset of ancient Pictish and present-day Scottish genomes, revealing partial population replacement taking place in eastern Scotland during the Anglo-Norman era (34). Morez et al. (2023) demonstrated that there was less genetic affinity between present-day eastern Scottish samples and the ancient Pictish genomes they sequenced, while present-day western Scottish samples shared significantly more genetic affinity with the ancient Pictish genomes (34). Furthermore, Morez et al. (2023) observed a clear genetic signature that differentiated the eastern Scottish population (showing a greater genetic affinity with populations with Anglo-Norman ancestry) from the western Scottish population (showing a greater genetic affinity with ancient samples of Iron-Age-British and Pictish ancestry) (34). Moreover, Gretzinger et al. (2022) observed a noticeable increase in Iron-Age-French ancestry among the present-day English population when compared to the ancient Iron-Age-British samples that have been sequenced (35). Gretzinger et al. (2022) concluded that Iron-Age-French-like ancestry was likely introduced to England during the early Medieval Age by people of Frankish-like origins and this continued through to the late Medieval Age with the Normans (35).

King and Jobling, (2009) argued that because of the correlation between surnames being traditionally inherited paternally, and Y-DNA being biologically passed on from father to son virtually unchanged, the study of Y-DNA has been of great value to genealogical studies investigating direct paternal ancestry (36). The consensus among researchers is that the comparison of mutations found within the Y-chromosome’s short tandem repeat markers (Y-STRs) and single nucleotide polymorphisms (Y-SNPs) between two or more males can accurately predict the degree of their patrilineal relationship, show how lineages evolved over time, and predict when lineages arose and diverged (37–41). Specific Y-SNP mutations on the Y-chromosome are also referred to as Y-DNA haplogroups and these define genetic population groupings (42). For example, the Y-SNP M417 is a mutation that defines the haplogroup R1a1a1, a major subclade (branch) of the haplogroup R1a (43, 44). The R1a-M417 haplogroup is especially significant because it represents the progenitor of a major expansion within the haplogroup R1a, and it is one of the main Y-DNA haplogroups associated with the spread of Indo-European languages (45). Genetic genealogical studies by Holton and Macdonald (2020), Stead (2023), Holton (2023), and DePew et al. (2024) have identified Y-SNPs that occurred in a specific individual or a narrow lineal group of patrilineal related individuals that lived within a known genealogical timeframe (46–49). A Y-SNP that can be specifically designated to an individual is termed a SNP-Progenitor (SNP-Progen) while Y-SNPs that can be identified as occurring in a lineal group of patrilineal related individuals are known as Multigenerational-SNPs (Multigen-SNPs) (46–48). The identification of these Y-SNPs is useful to genetic genealogists because they can confirm or reject hypotheses based on ambiguous documented evidence. Furthermore, once these Y-SNPs have been positively confirmed, they can be used to identify descendants that are not fortunate enough to be able to prove their descent with documentational evidence.

This study aimed to analyse Y-SNP data from FTDNA’s BigY700 tests to investigate whether the Y-DNA testing of proposed well-documented living patrilineal descendants of William de la HAYA II supports the genealogical consensus that his direct paternal lineage has survived to the present. The Y-SNP data from testing the BigY700 allows genetic genealogists to estimate a time to the most recent common ancestor (TMRCA) when two or more individuals share a specific Y-SNP, but their common ancestor is unknown. The mutation rates provide a date range for the occurrence of all Y-SNPs that are shared between two or more testers and if the common ancestor between two or more matching testers is known, their shared Y-SNPs can be assigned to specific individuals. Another aim is to identify SNP-Progens or Multigen-SNPs associated to the Hay noble lineage, so any EPP events can be identified using Y-DNA testing. Furthermore, the study aims to use the data for SNP-Progens or Multigen-SNPs to help identify Hay males of direct paternal descent who cannot prove their lineage due to gaps in the documentary records. The data from this study also have the potential to provide evidence to help reach a reliable conclusion on the proposed Norman origins of William de la HAYA I. Furthermore, the analysis of Y-DNA haplogroups found among Family Tree DNA testers with the surname Hay (plus associated spelling variants: Hays, Haye, Hayes, Hey and Haya) will be assessed to reach conclusions on the level of haplogroup diversity. An analysis into Y-DNA haplogroup diversity will provide data to evaluate the hypothesis made by Durie (2022) [20], who suggested that surnames in Scotland were largely adopted for many reasons, and that a surname of noble origin does not mean an individual is descended from the noble progenitors of that surname. Finally, this study will provide further insight into Y-SNP mutation rates using the BigY700 test results of three well-documented Hay patrilineal descendants who can trace their ancestral pedigree back to the 12th century.

## Methodology

Holton and Macdonald (2020) highlight the significant challenges faced when researching well-documented living descendants of medieval noble lineages (46). However, they confirmed that there are many examples of individuals of various ancestries who have the good fortune of being well documented, having extant lines of descent to the present day (46). Research by Larmuseau et al. (2014), Stead (2023), and DePew et al (2024) also demonstrates the existence of well-documented lineages back to the high to late Middle Ages (c. 1000-1500 CE) (47, 49, 50).

## Traditional genealogical documentary research

The methodology implemented in this study utilised a mixed disciplinary approach that combined traditional genealogical research, genetic genealogical research and archaeogenetic research methods. Holton and Macdonald (2020) demonstrated that authoritative documentary sources can be used to identify living descendants of noble lineages (46). Therefore, a range of publicly available authoritative documentation, primary and secondary sources (Table 1) were used to identify three potential test takers with proven patrilineal descent from William de la HAYA II, 1^st^ of Erroll.

**Table 1.**
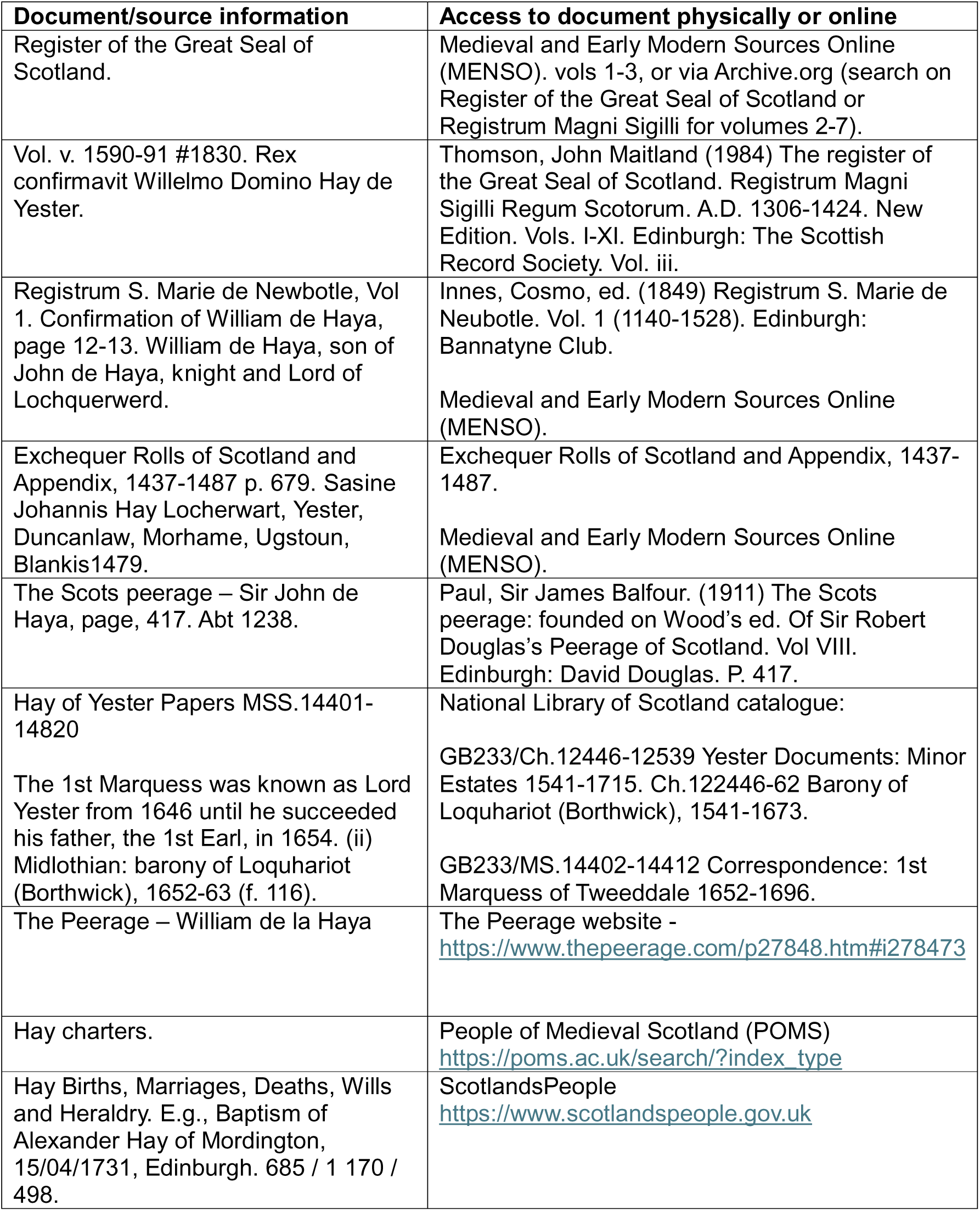
Examples of the sources consulted to prove the ancestries of the three test takers used for this study.

| Document/source information | Access to document physically or online |
| --- | --- |
| Register of the Great Seal of Scotland. | Medieval and Early Modern Sources Online (MENSO). vols 1-3, or via Archive.org (search on Register of the Great Seal of Scotland or Registrum Magni Sigilli for volumes 2-7). |
| Vol. v. 1590-91 #1830. Rex confirmavit Willelmo Domino Hay de Yester. | Thomson, John Maitland (1984) The register of the Great Seal of Scotland. Registrum Magni Sigilli Regum Scotorum. A.D. 1306-1424. New Edition. Vols. I-XI. Edinburgh: The Scottish Record Society. Vol. iii. |
| Registrum S. Marie de Newbotle, Vol 1. Confirmation of William de Haya, page 12-13. William de Haya, son of John de Haya, knight and Lord of Lochquerwerd. | Innes, Cosmo, ed. (1849) Registrum S. Marie de Neubotle. Vol. 1 (1140-1528). Edinburgh: Bannatyne Club.<br>Medieval and Early Modern Sources Online (MENSO). |
| Exchequer Rolls of Scotland and Appendix, 1437-1487 p. 679. Sasine Johannis Hay Locherwart, Yester, Duncanlaw, Morhame, Ugstoun, Blankis1479. | Exchequer Rolls of Scotland and Appendix, 1437-1487.<br>Medieval and Early Modern Sources Online (MENSO). |
| The Scots peerage – Sir John de Haya, page, 417. Abt 1238. | Paul, Sir James Balfour. (1911) The Scots peerage: founded on Wood's ed. Of Sir Robert Douglas's Peerage of Scotland. Vol VIII. Edinburgh: David Douglas. P. 417. |
| Hay of Yester Papers MSS.14401-14820<br>The 1st Marquess was known as Lord Yester from 1646 until he succeeded his father, the 1st Earl, in 1654. (ii) Midlothian: barony of Loquhariot (Borthwick), 1652-63 (f. 116). | National Library of Scotland catalogue:<br>GB233/Ch.12446-12539 Yester Documents: Minor Estates 1541-1715. Ch.122446-62 Barony of Loquhariot (Borthwick), 1541-1673.<br>GB233/MS.14402-14412 Correspondence: 1st Marquess of Tweeddale 1652-1696. |
| The Peerage – William de la Haya | The Peerage website -<br>https://www.thepeerage.com/p27848.htm#i278473 |
| Hay charters. | People of Medieval Scotland (POMS)<br>https://poms.ac.uk/search/?index_type |
| Hay Births, Marriages, Deaths, Wills and Heraldry. E.g., Baptism of Alexander Hay of Mordington, 15/04/1731, Edinburgh. 685 / 1 170 / 498. | ScotlandsPeople<br>https://www.scotlandspeople.gov.uk |

Based on the outcome of the traditional genealogy research, it was hypothesised that all three potential testers should share a Y-SNP dated to have occurred between 1000-1150 CE because their genealogies suggest that they descend from William de la HAYA II, 1^st^ of Erroll. However, the genealogies of two of the potential testers suggested that they shared a more recent ancestor (John HAY, 1^st^ of Tweeddale, 1595-1653 CE) when compared to the third tester; therefore, likely sharing at least one more Y-SNP in common that occurred between 1150-1595 CE. Furthermore, it is important to highlight that the two well-documented testers that descend from John HAY, 1^st^ of Tweeddale can be confidently documented back to Sir John HAY (1200-1262 CE) and he is hypothesised to be the grandson of William de la HAYA II via his son Robert (8, 51).

To confirm that the well-documented individuals were the direct paternal descendants of the noble Hay lineage of Scotland, Y-DNA testing was required to investigate whether the individuals shared Y-SNPs that could be dated to around the time-period of their proposed common noble Hay ancestor. The identified living descendants were invited to participate in this study after the receipt of signed consent with the recruitment period staring on the 6^th^ of February 2024 and ending on the 7^th^ of October 2024. Ethical approval for this research was obtained from the University of Strathclyde’s Ethics Committee, reference number: UEC23/94. After signed consent was received, a non-invasive BigY700 test kit was posted out to each participant for cheek cell sample collection using the cotton mouth swabs provided in the test kit. Once received, each participant completed the DNA sample collection and then posted the sample back to the FTDNA laboratory for sequencing. The BigY700 test by Family Tree DNA (FTDNA) was the test of choice for this project for several reasons (52). Firstly, FTDNA was chosen due to their large Y-haplotree, currently consisting of 81,000 branches, 716,000 variants, and 566,000 SNP-tested users in their Y-DNA database (53). Secondly, FTDNA has strict policies that ensure tester privacy and security of data (54); they also provide a ‘closed project’ option for researchers that ensures a secure platform to carry out data analysis.

## Y-DNA testing strategy

This research opted to use FTDNA’s BigY700 test which is a next-generation sequencing (NGS) test that covers an approximate 23 Mb of the total 57.2 Mb of the GRCh38 (HG38) reference Y-chromosome (less than 1% of the whole human genome) (55). The BigY700 targets all the Y-SNPs present in the SNP rich regions most relevant for genetic genealogical research with up to 70 reads per position covered; however, any Y-SNP that returns a minimum of 10 derived (positive) reads is reported (52). FTDNA’s age estimates for SNPs that feature within their Y-DNA phylogenetic tree are calculated using the methodology developed by McDonald (2021) (56). McDonald’s approach calculates coalescence ages (times to most recent common ancestor, or TMRCAs) using a new, probabilistic statistical model that includes Y-SNP, Y-STR and ancillary historical data (56). McDonald demonstrated that this methodology provided highly accurate estimates by using the Y-DNA data of well-documented direct patrilineal Royal Stewart testers (56).

In this study, the Y-SNP data of the three documented testers were compared against their specific genealogies to identify SNP-Progens or Multigen-SNPs. FTDNA’s age estimates for specific Y-SNPs were accepted as the most accurate estimates since they were calculated using McDonald’s methodology (52, 56). The example in Figure 1 demonstrates how a SNP-Progen and Multigen-SNPs are identified by combining the documented genealogical information with the Y-SNP data of four fictitious testers. When two or more Y-DNA testers have well-documented ancestry that triangulates back to a direct paternal ancestor living during the medieval era, Multigen-SNPs and SNP-Progens can also be identified with a high degree of confidence.

**Fig 1.**
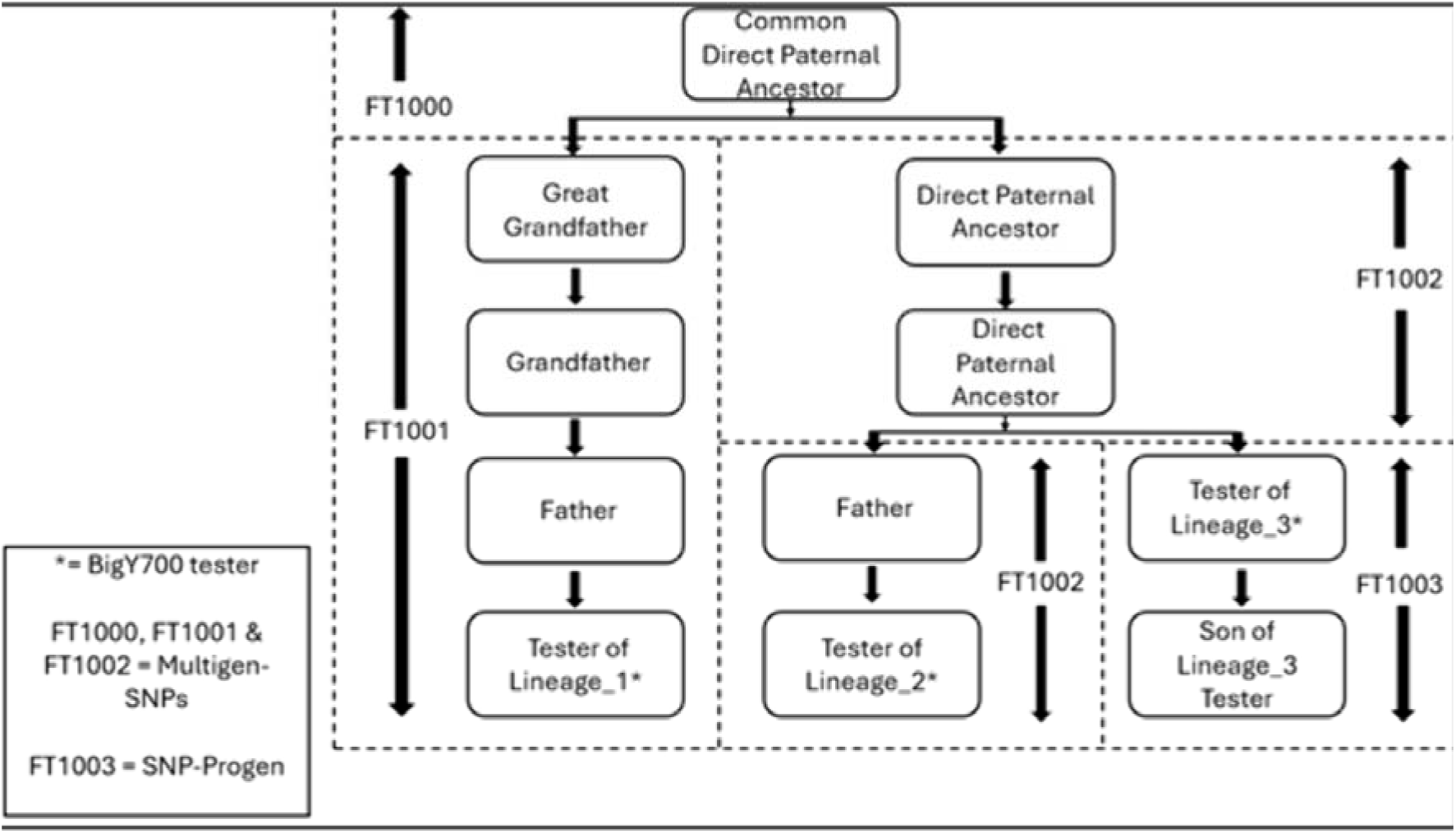
Diagram demonstrating how SNP-Progen and Multigen-SNPs are identified.

Figure 1 shows how three BigY700 testers are related to demonstrate how SNP-Progens and Multigen-SNPs are identified. Tester of Lineage_1* is the 3rd cousin of Tester of Lineage_2 via the same direct patrilineal line of descent, sharing the same 2nd great grandfather. Tester of Lineage_1 is the 2nd cousin, once removed of Tester of Lineage_3 and Tester of Lineage_3 is the uncle of Tester of Lineage_2. Since Tester of Lineage_1* is the only individual from his immediate patrilineal line, the Y-SNP FT1001 cannot be assigned to a specific individual; therefore, it is termed a Multigen-SNP as it could have occurred between Tester of Lineage_1*, his father, grandfather or great grandfather. Tester of Lineage_2 and Tester of Lineage_3 are more closely related, and they are derived (positive) for the Y-SNP FT1002; however, no new Y-SNPs have occurred in the father of Tester of Lineage_2 because Tester of Lineage_2 remains FT1002 which either occurred in his grandfather of great grandfather (another example of a Multigen-SNP). Furthermore, Figure 1 shows that Tester of Lineage_3 can be identified as the SNP-Progen because he is the brother of Tester of Lineage_2’s father and Lineage_3 carry an additional Y-SNP called FT1003; therefore, this Y-SNP must have occurred in Tester of Lineage_3, making him the SNP-Progen for FT1003. Based on the information shown in Figure 1, everyone in the phylogenetic tree will have their Y-SNP ancestral path written as shown in Table 2.

**Table 2.** Y-SNP ancestral path for the individuals shown in Figure 1.

| Individuals shown in Figure 1 | Y-SNP ancestral path |
| --- | --- |
| Common Direct Paternal Ancestor to all three lineages | FT1000 |
| Great Grandfather, Grandfather, Father and Tester of Lineage_1* | FT1000 > FT1001 |
| Great Grandfather, Grandfather and Father of Tester of Lineage_2* | FT1000 > FT1002 |
| Tester of Lineage_3* and his Son | FT1000 > FT1002 > FT1003 |

To investigate the degree of Y-DNA haplogroup diversity among individuals with the surname Hay, this study utilised the Y-DNA data of 109 testers that is publicly available on the Hay surname FTDNA project (57). In situations where a tester has not carried out any specific Y-SNP testing and they have only tested Y-STR markers, FTDNA only provide a very general Y-DNA haplogroup prediction. The STR data of the testers that had tested between 67-111 STR markers were inputted into the NEVGEN tool (n=29) (58) to predict a more refined Y-DNA haplogroup as implemented by Zhabagin et al (2024) (59). For example, Tester: 398281 was only assigned R1a-M512 by FTDNA; however, NEVGEN was able to predict a more refined Y-DNA haplogroup for tester using their 67 STR markers. NEVGEN accurately predicted kit number: 398281 as being positive for R1a-YP287 (58), a SNP several mutations further downstream of R1a-M512 as illustrated below: R1a - M207 > M173 > M420 > M459 > M512 > M417 > CTS4385 > Y2894 > L664 > FGC41399 > S2894 > YP287.

Once all the data were collated, testers were grouped based on the closest major Y-DNA haplogroup they were associated with. For example, kit number: 398281 was grouped in the R1a > L664 grouping.

## Results

### Y-DNA Haplogroup diversity among testers with the surname Hay (plus spelling variants)

The first set of results demonstrate the Y-DNA haplogroup diversity observed among the testers with the surname Hay (plus spelling variants) that are participating in the Hay FTDNA project. Figure 2 shows that Y-DNA haplogroups R1a, R1b, I1, I2, J1 and Q were all represented.

**Fig 2.**
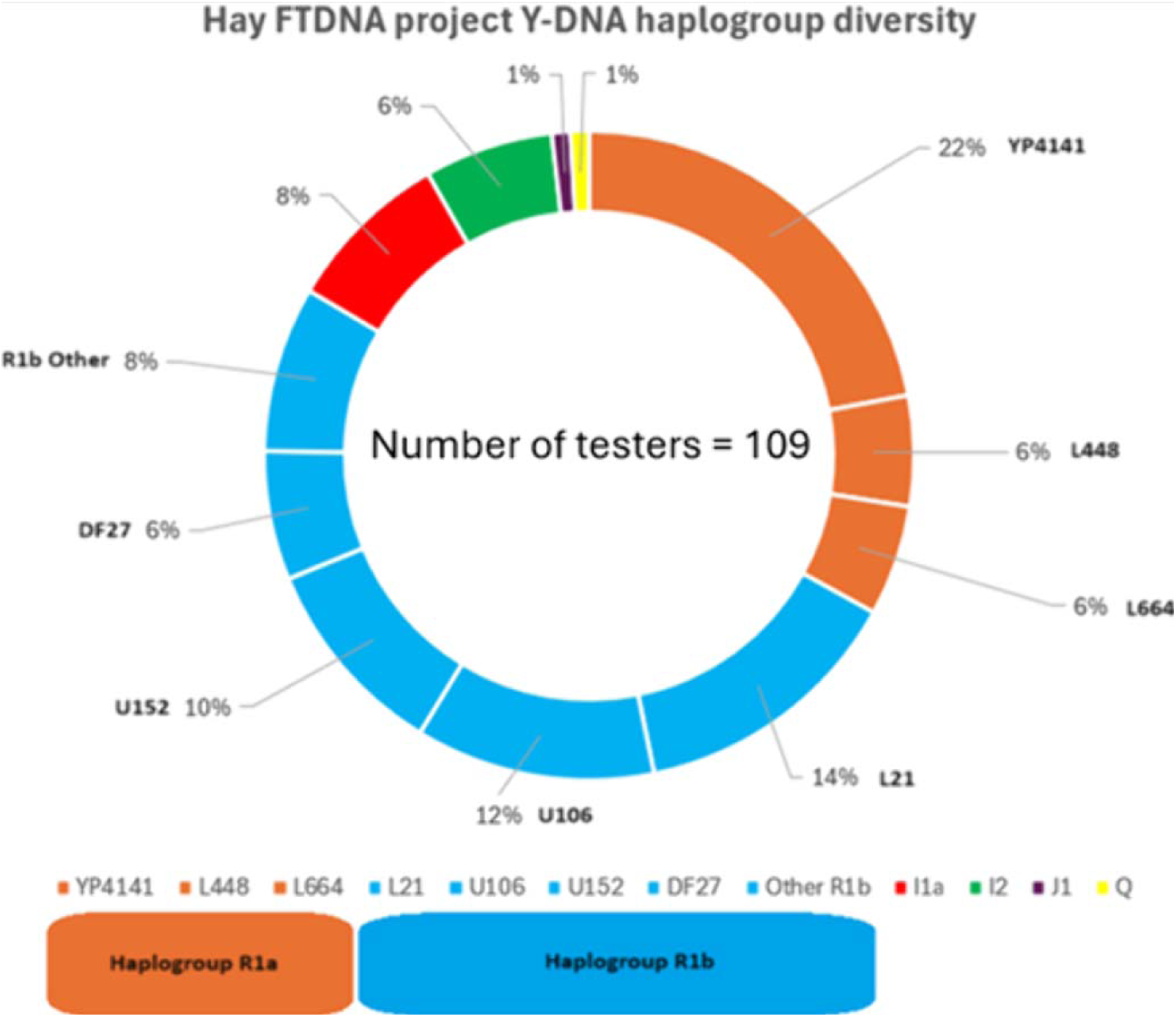
Y-DNA haplogroup diversity among testers with the surname Hay (plus associated spelling variants) within the Hay FTDNA project.

The evidence also demonstrates significant diversity within specific haplogroups (e.g., R1a > YP4141; R1a > L448 & R1a > L664) with haplogroup R1b proving to be the most diverse haplogroup represented. The Y-DNA diversity observed in this study is similar to previous research completed on Clan Forbes (47).

### The Y-DNA results of the three well-documented noble Hay descendants

The BigY700 test results of the three well-documented noble Hay descendants showed that they all shared the following Y-SNP ancestral path (see Table 3): R1a-M420 > YP4141 > YP4132 > YP4169 > YP4208 > YP4131 > YP4138 > YP6500 > FTT161.

**Table 3.** Basic Y-SNP ancestral path and specific Y-SNP details.

| **Y-SNP name** | **Location** | **Ancestral allele** | **Derived allele** | **FTDNA TMRCA estimate ranges (Earliest < Median > Latest; 95% probability)** |
| --- | --- | --- | --- | --- |
| FTT161 | CP086569.2: 12101060 | A | G | 877 CE < **1081 CE** > 1251 CE |
| YP6500 | CM000686.2: 7696598 | C | G | 687 CE < **930 CE** > 1131 CE |
| YP4138 | CM000686.2: 6863798 | G | T | 582 CE < **824 CE** > 1027 CE |
| YP4131 | CM000686.2: 2862540 | A | G | 129 BCE < **219 CE** > 513 CE |
| YP4208 | CM000686.2: 19384185 | C | A | 3795 BCE < **2945 BCE** > 2218 BCE |
| YP4169 | CM000686.2: 12489924 | T | C | 4193 BCE < **3291 BCE** > 2581 BCE |
| YP4132 | CM000686.2: 2880609 | C | G | 5481 BCE < **4406 BCE** > 3483 BCE |
| YP4141 | CM000686.2: 7404757 | C | T | 13078 BCE < **11025 BCE** > 9250 BCE |
| M420 | CM000686.2: 21311315 | T | A | 17707 BCE < **15272 BCE** > 13137 BCE |

Bonito et al (2021) suggested that the palindromic regions of the Y-chromosome are highly prone to gene conversion events; therefore, the identification of SNPs may be complex when performing short-read targeted NGS (60). Hallast et al (2023) identifies the four heterochromatic subregions in the human Y-chromosome: the (peri-)centromeric region, DYZ18, DYZ19 and Yq12 as highly repetitive (61); therefore, these regions are less suitable for the identification of reliable Y-SNPs using NGS. Figure 3 is a map of the Y-chromosome that identifies the positions of the Y-SNPs shown in Table 3. These Y-SNPs are located within the non-recombining portion of the Y-chromosome (NRY), where Y-SNPs occur at a relatively slow rate and without the instability caused by gene conversion, deletions, and duplications. Furthermore, these Y-SNPs are not located in any of the regions that are deemed unreliable for Y-SNP calling; therefore, these Y-SNPs are regarded to be highly stable over generations, making them reliable for phylogenetic studies.

**Fig 3.**
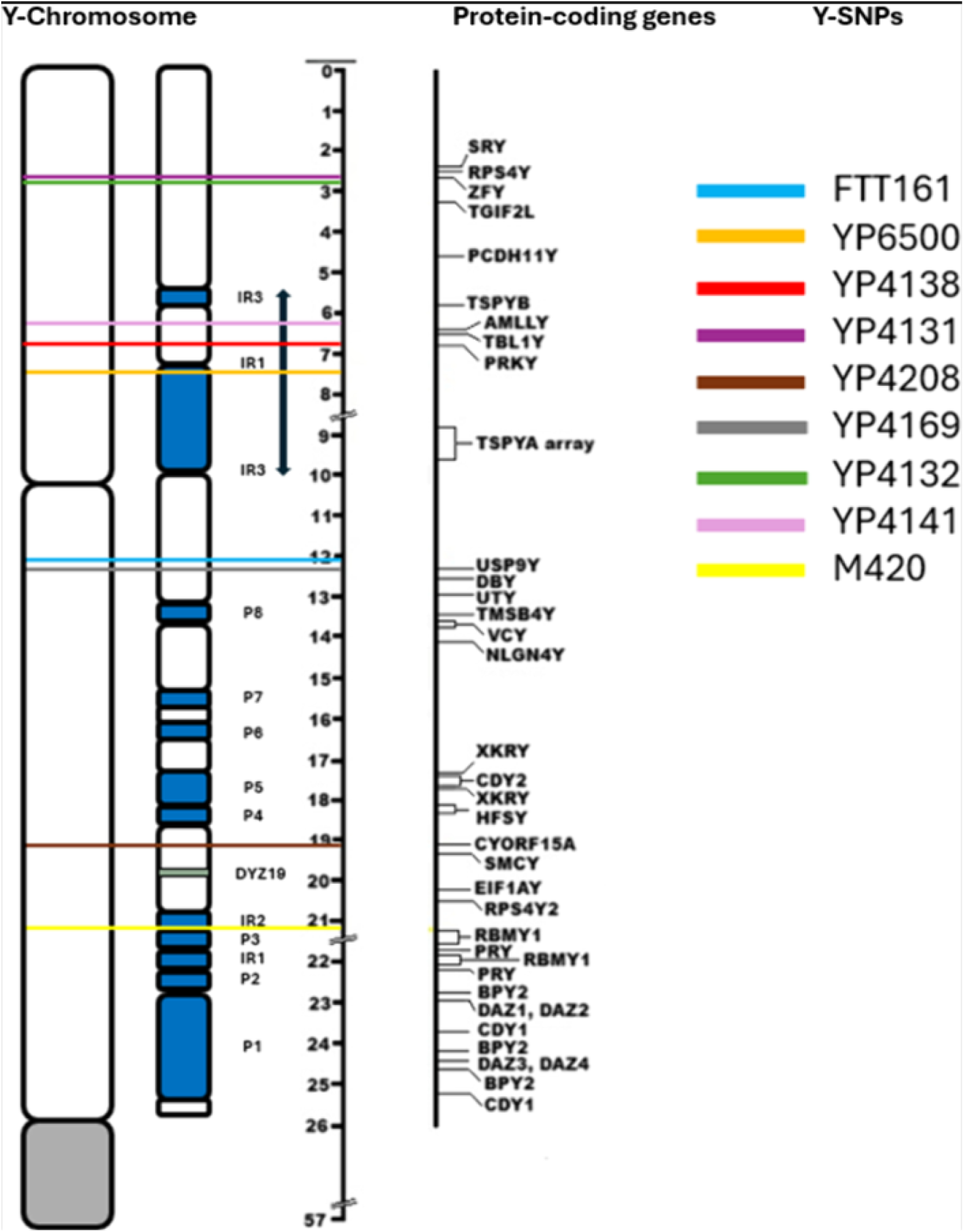
Y-Chromosome map adapted from Jobling and Tyler-Smith (2003) showing Y-SNPs associated with the noble Hay lineage (62)

The sequences in Table 4 show the bases surrounding the Y-SNPs associated to the ancestral path for the noble Hay lineage. It can be observed that each derived Y-SNP is surrounded by non-repetitive sequences; therefore, increasing the likelihood that these Y-SNPs are stable and phylogenetically relevant.

**Table 4.**
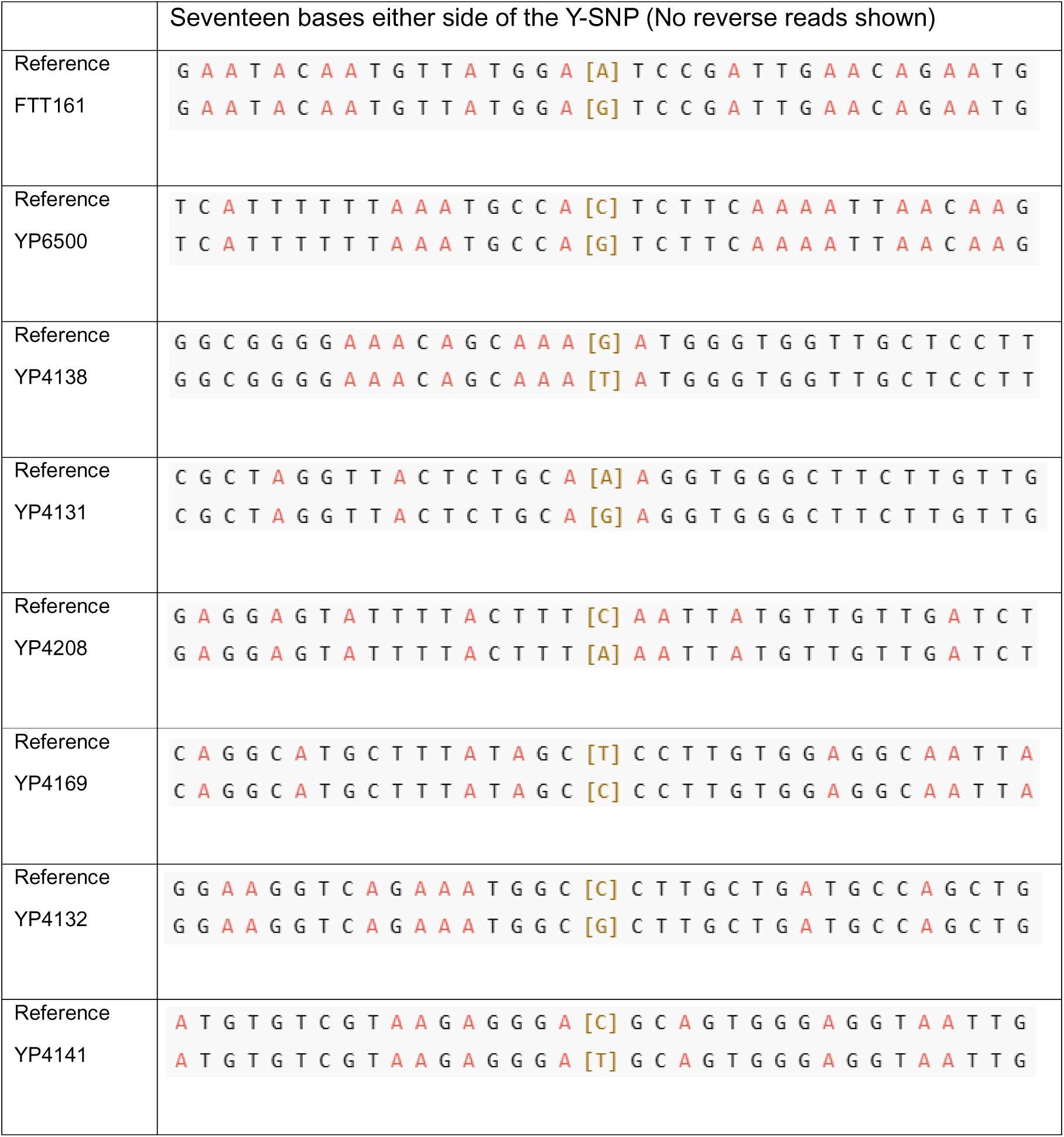
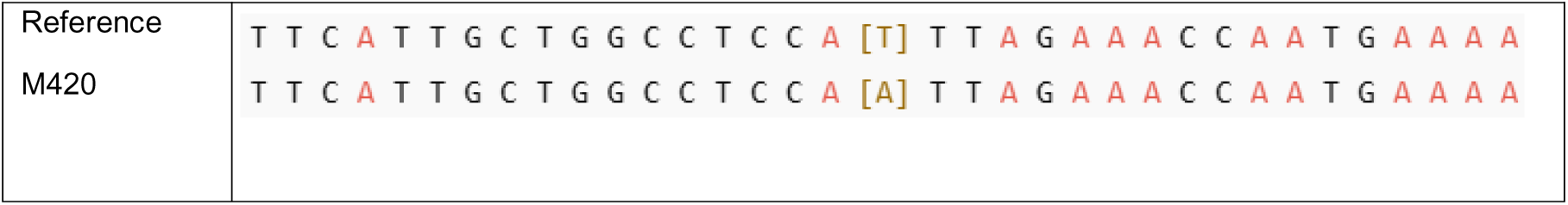
Sequencing data of the bases surrounding the Y-SNPs identified in the Y-SNP pathway for the Hay noble lineage.

The first line in each row of Table 4 shows the 17 bases of sequence surrounding either side of each Y-SNP found in the Y-SNP pathway for the Hay noble lineage. The sequences are in the Genome Reference Consortium Human Build 38 (GRCh38) format. The 18^th^ base in each sequence is coloured brown and boxed using square brackets to indicate the location of the specific SNP (e.g., FTT161 = [A] ancestral allele; [G] derived allele).

The Y-SNP FTT162 is phylogentically positioned downstream of FTT161 and it defines a noble Hay branch that has a significant number of descendants; n=47 testers at the YFull database (63). This Y-SNP is located in the highly repetitive heterochromatic region of the Y-chromosome (see Figure 4). This region is regarded as having large portions of noncoding DNA that is typically prone to mutations caused by replication slippage (64–66); therefore, Y-SNPs occurring within this region of the Y-chromsome need to be treated with caution as they may not be phylogenetically relevant (67).

**Fig. 4.**
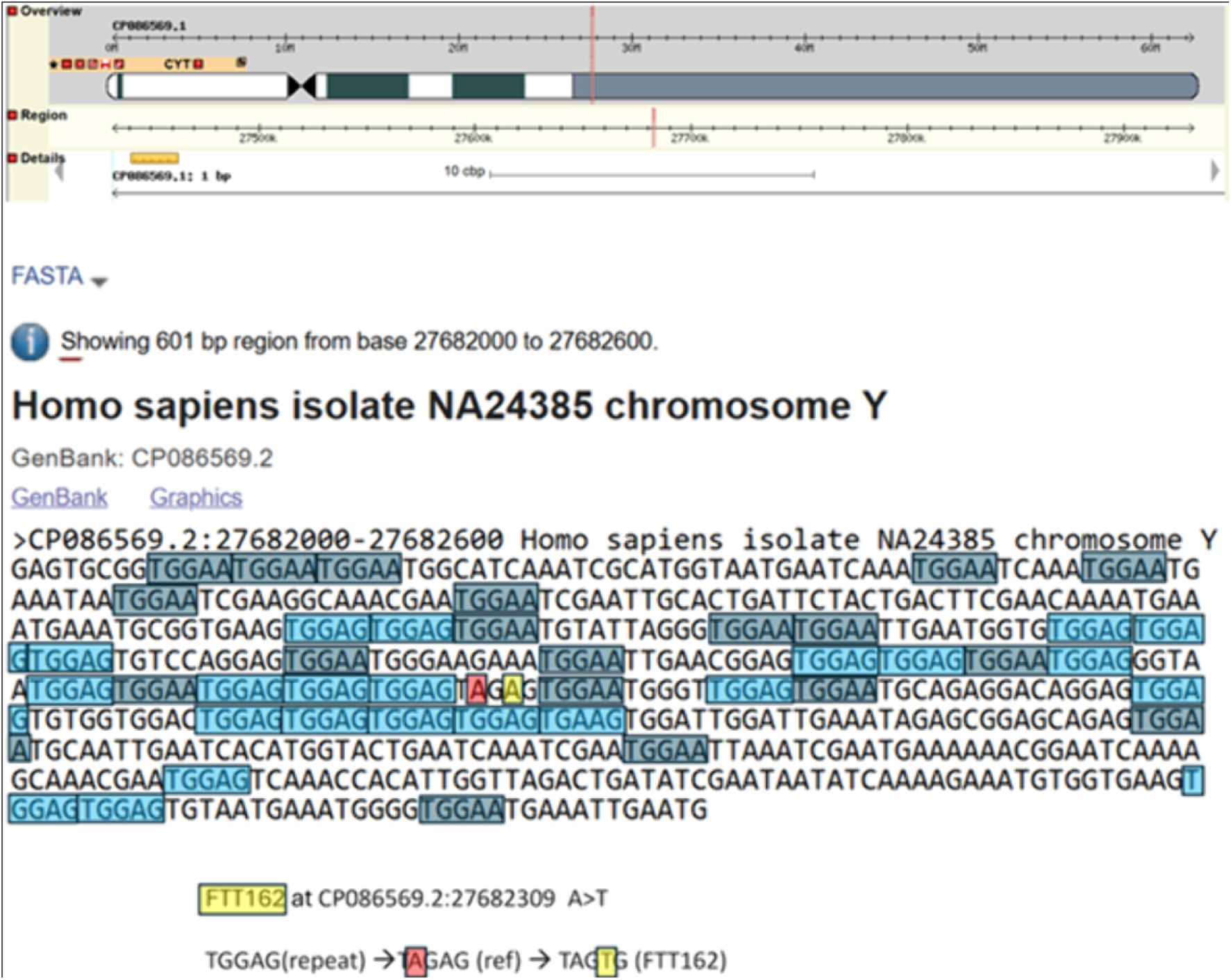
Y-SNP FTT162 location: CP086569.2:27682309: A > T and surrounding Sequence (68, 69)

Since FTT162 is located in a less stable region of the Y-chromosome, we analysed the surrounding sequence either side of this SNP to assess its stability and reliability of being phylogenetically relevant. Figure 4 highlights the repetitive sequences across a 600 bp region with FTT162 in the middle of the sequence and annotated in yellow. The way that FTT162 likely formed is by a change from the TGGAG repeat to TAGAG (TGGAG --> TAGAG; G>A; annotated in red), and this terminated the TGGAG repeated sequence that resulted in stabilising this section of sequence. After the TAGAG section had changed, the resulting stability provided a unique background for the FTT162 SNP (A>T; annotated in yellow) to occur. When FTT162 occurred, it changed the dynamics of the surrounding repeats so that two out of five bases were different to the left and to the right of the repeat units either side of FTT162. Therefore, these changes would have increased the stability at FTT162, making it an unlikely place for the DNA polymerase to slip, thus, restricting any conversion to either of the neighbouring repeats; TGGAG or TGGAA, respectively. For the reason discussed above, we have no concerns on the stability and reliability of this Y-SNP.

The three well-documented Hay testers also match a significant number of testers (n=24/109 at the Hay FTDNA project) with the surname Hay (plus associated spelling variants) who also tested positive for at least FTT161 (70). At this point, it is unclear whether YP6500 (plus equivalent SNPs BY33394 / FT2017) are Y-SNPs specific to the Hay noble lineage or whether they occurred in an ancestor that predates William de la HAYA, I of Normandy.

Furthermore, the Y-SNPs BY33394 and FT2017 are assumed to be equivalent to YP6500 as it is unclear whether they appeared simultaneously in the evolutionary timeline, or whether their occurrence was totally independent of each other. New data often changes the status of assumed equivalent Y-SNPs by proving that they occurred independently of other Y**-**SNPs. Therefore, it is never a guarantee that any Y-SNP is equivalent to other Y**-**SNPs. Once new data provides enough evidence to show that assumed equivalent Y-SNPs did occur independently, their position within the phylogenetic tree changes accordingly to represent their correct chronological order. The results show that there are some non-Hay surnames among the testers (n=12) who are positive for FTT161 and downstream SNPs, these non-Hay surnames are likely the result of an EPP taking place. The results strongly suggest that FTT161 is a Y-SNP that defines the noble Hay lineage with the downstream Y-SNPs of FTT161 identifying descent from William II de la HAYA, 1^st^ of Erroll, (d. 1201). Most testers with the surname Hay on the Hay FTDNA project (n=85/109) are not direct paternal descendants of the Hay noble progenitor (70). Figure 5 shows the phylogenetic block-tree of R1a-YP4138 which splits to form the subclades YP6500 (ancestor of the Hay noble lineage) and three parallel subclades, CTS11317, BY190800 and BY234055, respectively. The subclades CTS11317, BY190800 and BY234055 have several testers with surnames of Norman origin (Manwarren/Manering, Pickering and Travers) (71), and several testers with surnames of English origin. The non-Hay surnames associated to the parallel subclades of YP6500 is expected since the age estimate for R1a > YP4138 is 582 < 800 > 1027 CE (95% probability) (72); a timeframe generally accepted as being before the earliest known adoption of surnames in Europe (73).

**Fig 5.**
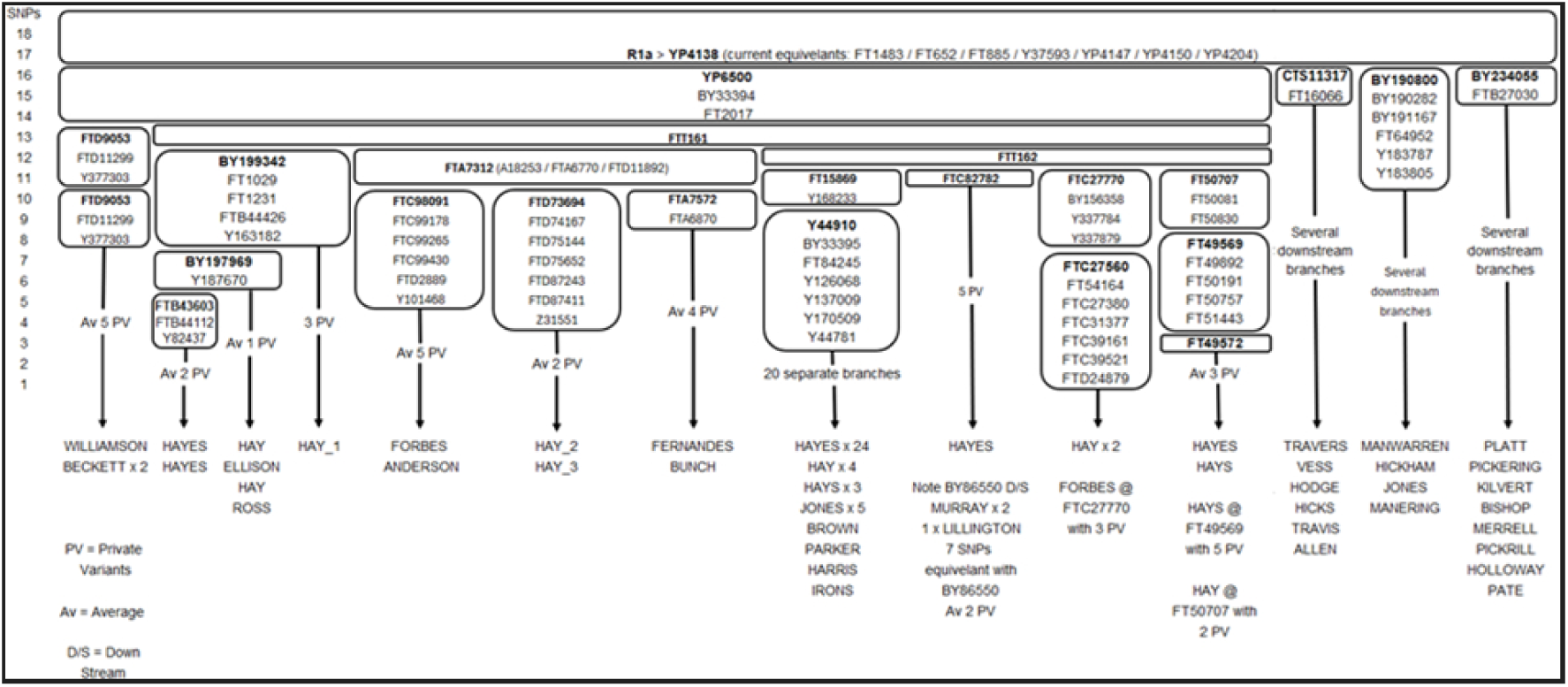
R1a > YP4138 Phylogenetic Block-Tree showing the Y-SNPs associated to the descendants of the Hay noble progenitor YP6500.

Figure 6 was produced by combining the Y-SNP data with the ancestral pedigrees of the three well-documented Hay testers.

**Fig 6.**
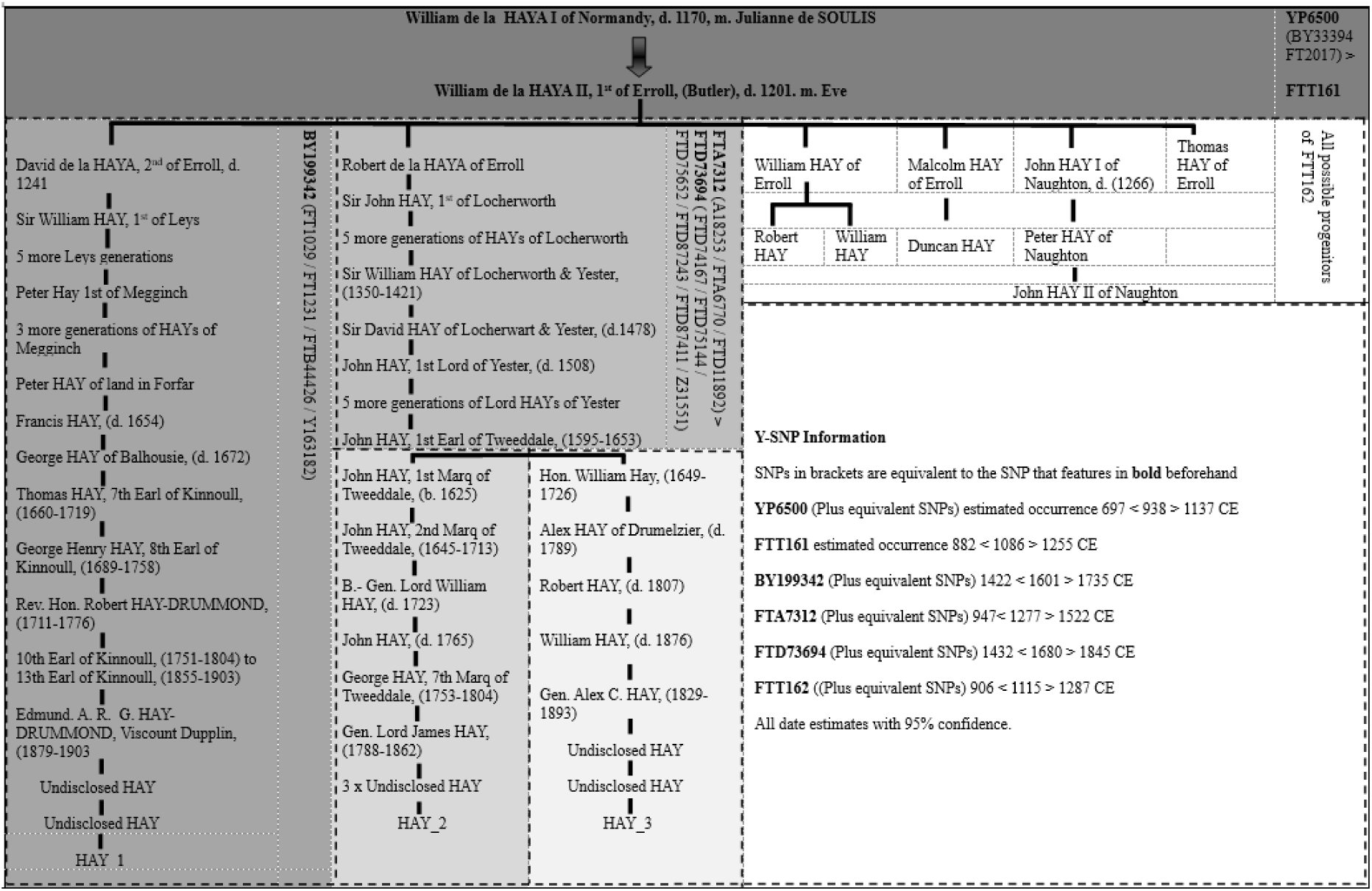
Original Hay Noble phylogenetic tree using the pedigrees of the three well-documented noble Hay descendants.

The phylogenetic tree identifies several Multigen-SNPs associated to the noble Hay lineage. Testers HAY_1, HAY_2 and HAY_3 are the three well documented individuals who were identified during the traditional genealogical research phase of the study. The other testers that match the three well-documented Hay individuals can also be placed in approximate positions within the Hay noble phylogenetic tree based on their Y-SNPs; however, further documentational evidence or the testing of more well-documented descendants will be required for a more accurate positioning. Three clear subclades that are phylogenetically downstream of the Y-SNP FTT161 were identified in this study (BY199342, FTA7312 & FTT162). The combined Y-SNP and ancestral documentation data suggests lines of descent from three of the six sons of William II de la HAYA, 1^st^ of Erroll. The Y-SNP BY199342 (plus equivalents) that were found in tester HAY_1 can confidently be assigned to the descendants of David de la HAYA, 2^nd^ of Erroll (d. 1241 CE), first son of William II de la HAYA, 1^st^ of Erroll (d.1201 CE). However, it is uncertain whether BY199342 occurred in David de la HAYA, 2^nd^ of Erroll or in one of his immediate direct descendants. Furthermore, HAY_2 and HAY_3 are negative for BY199342 (plus, equivalents) and positive for FTA7312 (plus equivalents) while likely descending from Robert de la HAYA of Erroll (the younger brother of David de la HAYA, 2^nd^ of Erroll). The Y-SNP data and the pedigrees of the well-documented testers show a clear genetic divergence that was previously hypothesised.

Moreover, the results support the documentary evidence that suggests that HAY_1, HAY_2 and HAY_3 are descendants of William II de la HAYA, 1^st^ of Erroll with a high degree of certainty.

### Analysis of the mutation rates observed among the well-documented Hay testers

The results in Table 5 suggest that the Y-SNP mutation rate observed in the Hay noble lineage does not fit the standard average mutation rate proposed by McDonald, (2021); however, the mutation rates do fall within the accepted mutation rate ranges. For example, FTD73694 is dated to 1700 CE with a date ranging from 1428 to 1844 CE with 95% probability; however, the TMRCA has been identified to 1595 CE with 100% confidence (56). McDonald proposes that the standard molecular clock for Y-SNP mutation rate average is 83 years/Y-SNP mutation (56). The data from the three well-documented Hay testers show an average mutation rate of 67.8 years/Y-SNP between 1140-1950 CE. The results for HAY_2 and HAY_3 also show an average mutation rate of 41.4 years/Y-SNP between 1140-1595 CE.

**Table 5.** Y-SNP analysis of the three well-document Hay testers.

| Participant cohort | Total number of Y-SNPs downstream of FTT161 | Number of generations since William HAY, 1st Earl of Erroll, b. 1140 CE | Average years per generation | Average Y-SNP mutation rate (yrs) |
| --- | --- | --- | --- | --- |
| HAY_1 | 11 | 25 | 32.4 | 73.6 |
| HAY_2 | 12 | 26 | 31.2 | 67.5 |
| HAY_3 | 13 | 24 | 33.75 | 63.3 |
| **Total average from HAY_1, HAY_2 & HAY_3 bet. 1140 – 1950 CE** | **12** | **25** | **32.45** | **67.8** |
| **HAY_2 & HAY_3 bet. 1140-1595 CE** | **11** | **16** | **28.4** | **41.4** |

Additionally, only one private variant (unique Y-SNP) occurred in HAY_2, and only two private variants occurred in HAY_3 since their common ancestor John HAY, 1^st^ Earl of Tweeddale (1595–1653). Figure 7 illustrates the average mutation rates observed between HAY_2 and HAY_3.

**Fig 7.**
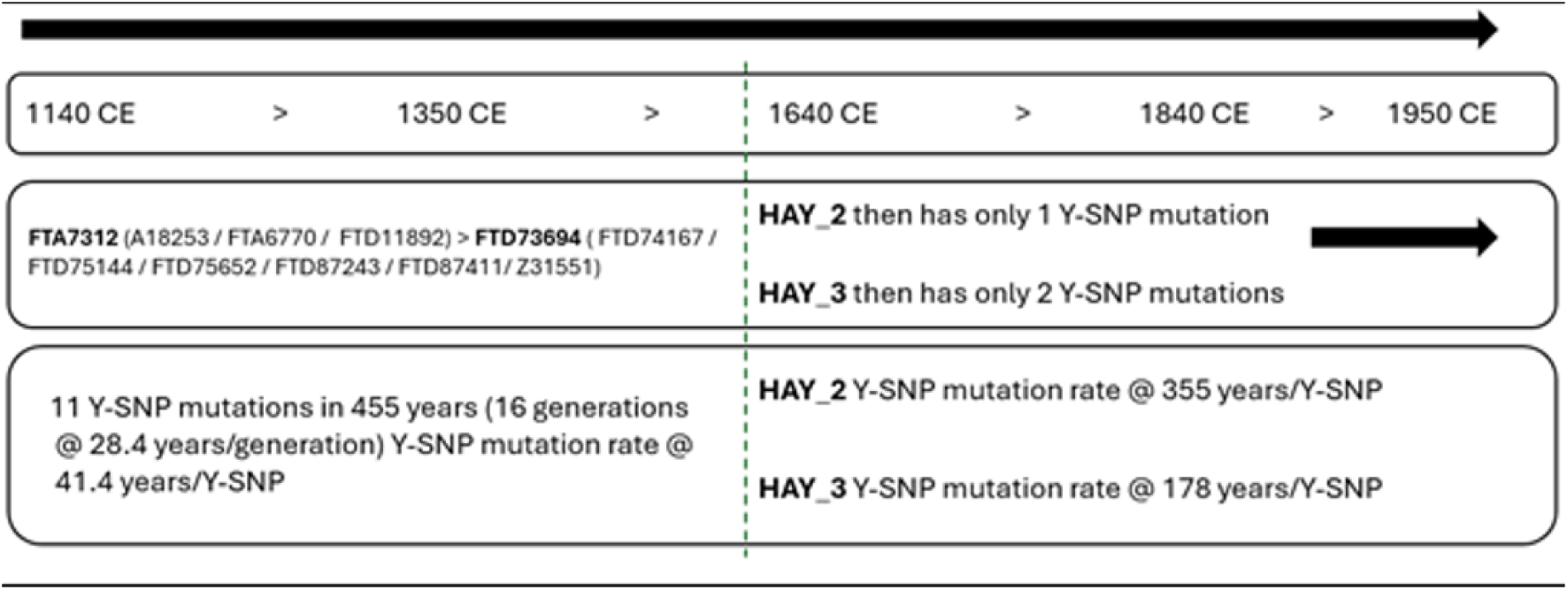
Diagram showing average mutation rate of Y-SNPs for HAY_2 and HAY_3.

The observation of variation in mutation rates during specific time periods could explain a proposed documented pedigree for an existing FTDNA tester with the surname HAY, who is positive for the subclade FTT162 (referred to as HAY_4 in this report). The documentary evidence for HAY_4 suggests possible descent from Robert de la HAYA of Erroll, younger brother of David de la HAYA, 2^nd^ of Erroll. This proposed ancestry would need to span approximately two hundred years, totalling eight generations between William de la HAYA II, 1^st^ of Erroll and Sir William HAY of Locherworth and Yester, Sheriff of Peebles, (1350-1421 CE) which can be seen in Figure 8.

**Fig 8.**
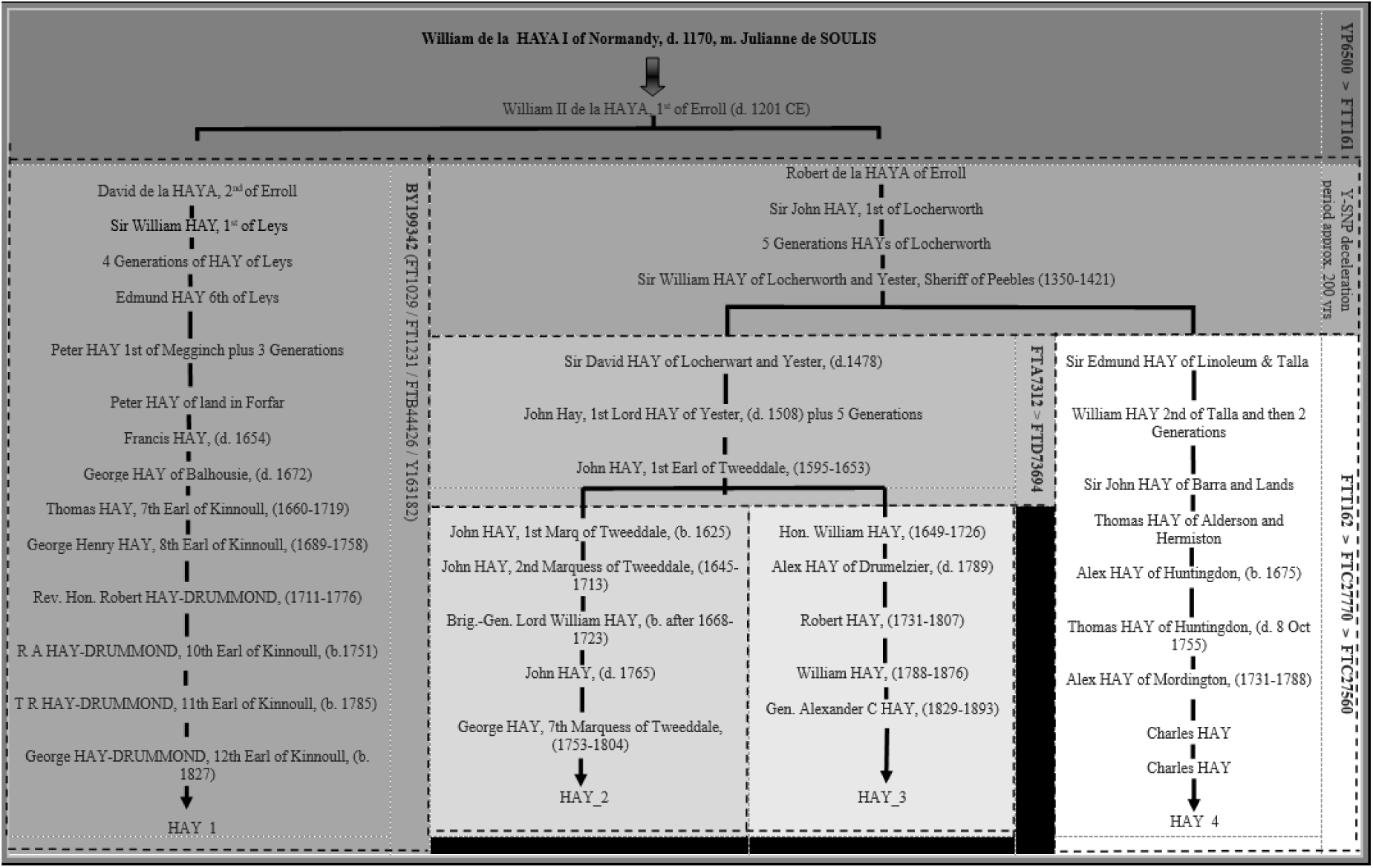
Alternative Hay Noble phylogenetic tree when HAY-4’s proposed pedigree is considered.

If HAY_4’s pedigree is correct, the eleven Y-SNPs shared between HAY_2 and HAY_3 would have to have occurred during a period between 1350-1595 CE (245 years). In addition, Hay_4’s pedigree would require an average Y-SNP mutation rate of 1 x Y-SNP/22.3 years with an average generation rate of 30.6 years (Figure 9).

**Fig 9.**
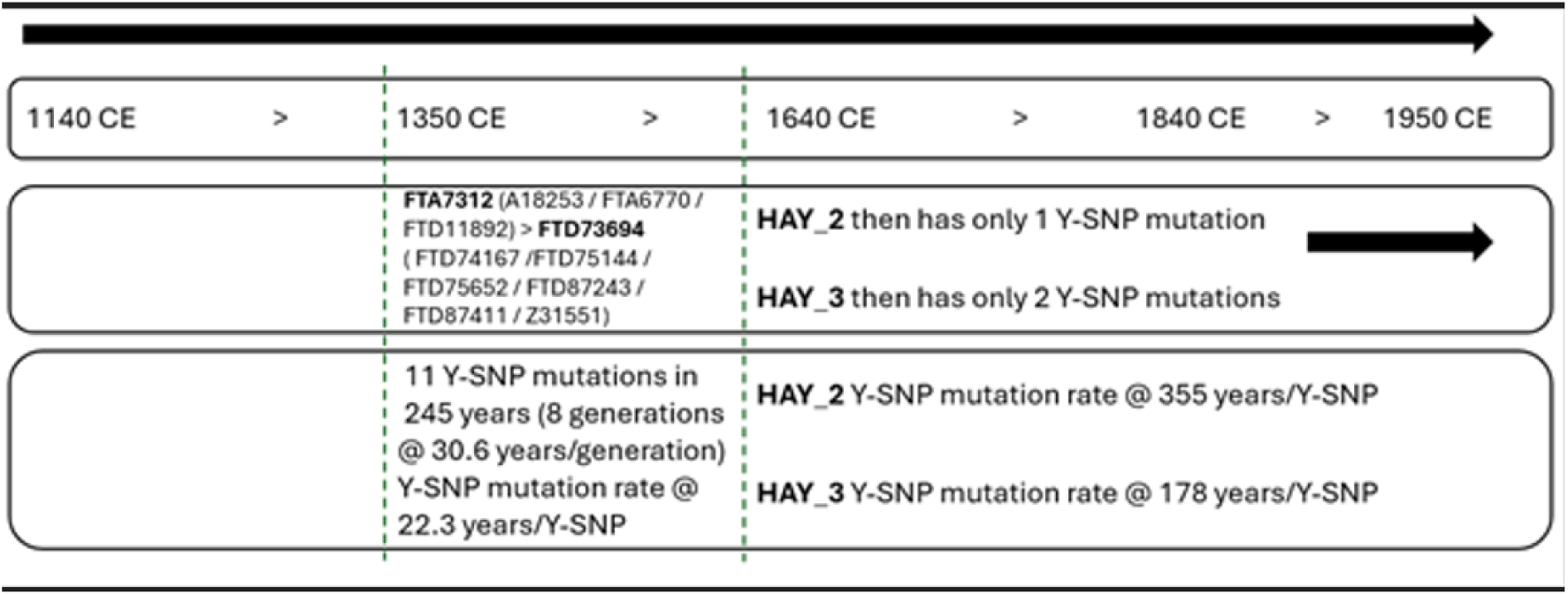
Diagram showing the average Y-SNP mutation rate if HAY_4’s proposed pedigree is considered.

The average mutation rate of 22.3 years/Y-SNP that is associated to HAY_4’s proposed pedigree falls within the possible mutation rate range since FTT162 is estimated to have occurred in 1107 CE, with a range between 898 CE and 1280 CE with 95% probability (74). Therefore, this ancestral pedigree of HAY_4 is possible on a genetic level. However, there are alternative ancestral pedigrees for HAY_4, but these cannot be documented back to William II de la HAYA, 1^st^ of Erroll, (d.1201) due to a lack of records. To validate HAY_4’s true line of descent back to William II de la HAYA, 1^st^ of Erroll, the Y-DNA testing of a well-documented direct paternal descendant of the four remaining sons of William II de la HAYA, 1^st^ of Erroll (William, Malcolm, Thomas and John HAY I of Naughton) would be required.

## Discussion

The Y-DNA haplogroup diversity observed among the Hay FTDNA project testers demonstrate a lack of homogeneity among the FTDNA testers with the surname Hay (plus associated spelling variants). Only 22% of the FTDNA testers with the surname Hay descend from a Hay nobleman, defined by the Y-SNPs YP6500 and FTT161 (plus subclades). A similar percentage of Y-DNA haplogroup diversity was also observed in a Clan Forbes genetic study where Stead (2023) reported 27% of FTDNA testers within the Clan Forbes project were shown to be direct paternal descendants of the noble lineage (47).

Furthermore, the results also support the hypothesis made by Durie (2022) who suggested that surnames in Scotland were largely adopted for many reasons, and that a surname of noble origin does not mean an individual is descended from the noble progenitors of that surname (23). More research needs to be conducted on other surnames associated to Scottish nobility for comparison to this study and the Clan Forbes genetic study (47). With further data, more specific questions can be asked; for example, are there specific historical events that facilitated surname changes, and can this be evident in Y-DNA data?

The Y-SNP YP4138, plus equivalents, are directly upstream of YP6500 in the phylogenetic tree. Subclades of YP4138 are found among several testers with the Norman-in-origin surnames Manwarren, Pickering and Travers. therefore, strongly supporting the Norman origins of YP4138 and its subclades. The age estimate for YP4138 is 582 < 800 > 1027 CE (95% probability) (72), therefore, also aligning with the Norman origin hypothesis. Some testers with the surname Travers have well-documented ancestry from the Travers noble lineage of Horton, Cheshire (75) and Cork, Ireland (76). Although the Travers lineage of Horton and Cork can only be confidently documented back to the mid-15th century, they could be descended from the Norman ancestor of Ralph Travers of Berney, c. 1197 (75, 76). Further testing of well-documented individuals with surnames of Norman origin that match the noble Hay and Travers testers could provide important Y-SNP data to help shed further light on their ancestral connections.

The research conducted in this study has provided strong evidence that the Multigen-SNPs YP6500 (plus equivalents) and FTT161 are key Y-SNPs that identify descent from the Hay noble lineage. Furthermore, two subclades of FTT161 (BY199342 & FTA7312) are Multigen-SNPs that indicate direct paternal descent from David de la HAYA, 2^nd^ of Erroll and his younger brother Robert de la HAYA of Erroll. Any individual that matches these key Y-SNPs can be confirmed as a direct paternal descendent of the Hay noble lineage with a high degree of confidence, even if documentary evidence is absent. Furthermore, the Y-SNP data can also provide more specific ancestral designations (e.g., descent from David de la HAYA, 2^nd^ of Erroll or his younger brother Robert). This dataset will become more refined when additional well-documented Hay testers complete BigY700 testing, leading to more Multigen_SNPs being assigned to specific Hay noblemen, and potentially identifying specific SNP-Progens. Invariant mutation rates across the human Y-chromosome have generally been accepted among researchers such as Xue et al, (2009), Helgason et al, (2015) and Kutanan et al, (2019) (77–79). The consensus among these authors were that the Y-SNP mutation rate in human Y lineages remained homogeneous; however, this was not supported by the research conducted by Ding et al, (2021) (80). Ding et al, (2021) suggests variable mutation rates across the human Y-chromosome haplogroups; however, it is not clear whether this heterogeneity is caused by interhaplogroup mutation rate variation (80). The Y-SNP data in this study supports the theory of variable mutation rates across the human Y-chromosome haplogroups (mean Y-SNP mutation rate of 67.8 years/SNP as opposed to the standard 83 years/SNP rate) (56).

This study is the first to observe the Y-SNP mutation rates of well-documented testers with ancestry tracing back to a common ancestor living in the 12th century. Although the data clearly demonstrate varying mutations rates during different time periods, it is unclear what causes this phenomenon. One explanation for this phenomenon could be that greater numbers of male offspring were born during the period where more Y-SNPs mutations are being observed. Furthermore, factors such as higher than average conception age of the Y-SNP progenitors could result in more Y-SNPs occurring, limitations with NGS, and specific haplogroup genetic mechanisms could also be at play.

The technological and methodological advances made by the telomere-to-telomere (T2T) consortium has already had a huge impact on identifying new Y-SNPs that were not reported with NGS technology (81). Although the BigY700 has identified several T2T Y-SNPs (e.g., FTT161 and FTT162) within the noble Hay phylogenetic tree (82), these Y-SNPs did not get recognised until T2T research was made available to researchers. Therefore, it is recommended that several noble Hay testers are eventually sequenced using the longread sequencing methodology implemented by the T2T consortium as the current research suggests that a higher resolution of the Y-chromosome can be achieved (81, 83, 84).

Furthermore, longread sequencing will likely develop the noble Hay phylogenetic tree by identifying new Y-SNPs, resulting in more accurate Y-SNP dating.

To the best of our knowledge, this study is the first to suggest variation in Y-SNP mutation rates using well-documented testers with proven pedigrees dating back to the 12th century.

To explore patterns in the variation of average Y-SNP mutation rates further, investigations into a wider range of well-documented lineages is recommended for future research. If similar variation of average Y-SNP mutation rates can be observed in other well-documented lineages, further research could focus on investigating possible causes for such variation, and the potential implications on Y-SNP dating. The dating estimates for Y-SNPs are very ambiguous due to the very large date ranges currently available. However, the identification of Multigen-SNPs and SNP-Progens provides far more precise dating of Y-SNPs, but the methodology is limited to well-documented lineages that can be traced back to the high Middle Ages era. Those lineages of noble descent are the only realistic candidates that can be reliably investigated to identify Multigen-SNPs and SNP-Progens back to the medieval era. Further limitations occur when a noble lineage has died out or researchers fail to recruit a well-documented individual for Y-DNA testing. Furthermore, investigating other noble lineages will provide more comparative data to permit conclusions on whether Y-DNA haplogroup diversity can be observed in other surnames associated to nobility. Y-DNA haplogroup diversity is likely facilitated by several mechanisms; however, specific historical events could be a key factor that facilitate the adoption of certain surnames. The impact of wars and migrations could be two key facilitators to explore in future research.

## Conclusions

The identification of Multigen-SNPs and SNP-Progens are of huge value to genealogical researchers. As demonstrated in this research and in other studies, Y-DNA testing has been proven to bridge significant gaps that exist in the documentary evidence when researching ancestral pedigrees (46–49). Furthermore, direct paternal descent that is proven by documentation is never a guarantee of true descent due to the probability of an EPP taking place. In reverse, not having a noble surname does not mean that an individual is not a direct paternal descendant of a noble lineage. The Hay testers had genetic matches with 24 individuals who have the surname Hay (plus associated spelling variants) and 12 individuals who did not have the surname Hay. Our genetic data proves that the 12 non-Hay testers are descendants of a noble Hay ancestor, and a surname change has occurred at some point in their ancestry, likely the result of an illegitimate conception sometime in the past. It is likely that several of the testers with the surname Hay are descended from a younger son of a noble lineage and these lineages eventually disappeared from the records as their descendants were landless. Research that focuses on Multigen-SNPs and SNP-Progen identification adds another dimension for validating pedigrees with a high degree of certainty. Furthermore, this type of research provides reliable data that can be used by researchers to challenge poorly researched pedigrees that make erroneous conclusions.

By comparing the Y-SNPs and ancestries of the three well-documented Hay noble descendants, we have managed to confidently identify several Multigen-SNPs associated to the noble Hay lineage. The genetic data for this study also provides further evidence that the Hay noble lineage originated in Normandy as they share the Y-SNP R1a-YP4138 with several individuals who have a surname of Norman origin. One specific tester that matches the Hay noble lineage prior to the birth of the noble Hay progenitor is proven to be a well-documented descendant of the noble Travers of Horton lineage. This research demonstrates that significant Y-DNA haplogroup diversity exists among FTDNA testers with the surname Hay. The Y-DNA haplogroup diversity observed in this study is strong evidence that supports Durie (2022) who suggested that most men with a surname associated to nobility will not be a direct paternal descendant of the noble progenitor (23).

## Acknowledgements

We gratefully acknowledge all sample donors who participated in this study. Thanks to Michael Travers, David Jones, Alan Hay, and the three well-documented Hay testers for their support. Thank you to Dr. Tunde Huszar and Graham S Holton for their support and expert input.

